# GAPS: Geometric Attention-based Networks for Peptide Binding Sites Identification by the Transfer Learning Approach

**DOI:** 10.1101/2023.12.26.573336

**Authors:** Cheng Zhu, Chengyun Zhang, Tianfeng Shang, Chenhao Zhang, Silong Zhai, Zhihao Su, Hongliang Duan

## Abstract

The identification of protein-peptide binding sites significantly advances our understanding of their interaction. Recent advancements in deep learning have profoundly transformed the prediction of protein-peptide binding sites. In this work, we describe the Geometric Attention-based networks for Peptide binding Sites identification (GAPS). The GAPS constructs atom representations using geometric feature engineering and employs various attention mechanisms to update pertinent biological features. In addition, the transfer learning strategy is implemented for leveraging the pre-trained protein-protein binding sites information to enhance training of the protein-peptide binding sites recognition, taking into account the similarity of proteins and peptides. Consequently, GAPS demonstrates state-of-the-art (SOTA) performance in this task. Our model also exhibits exceptional performance across several expanded experiments including predicting the apo protein-peptide, the protein-cyclic peptide, and the predicted protein-peptide binding sites. Overall, the GAPS is a powerful, versatile, stable method suitable for diverse binding site predictions.

## 1. Introduction

Peptides, built from sequences of different amino acids^1^, play key roles in many cellular processes within the organism^2^, including signal transduction cascades^3^, antimicrobial defense^4^, and immune responses^5^. Within the human proteome, more than 10^6^ peptide motifs are encoded^6^, and their functions are intrinsically related to interactions with target proteins^7^. To further the fundamental understanding of how proteins interact with peptides and to accelerate drug discovery development, an important challenge is to precisely identify the protein-peptide binding sites. However, experimental observation of these binding sites is laborious and time-consuming^7^. Recently, technology development has promised to provide a better grasp on computational predicting protein-peptide binding sites, and many methods are proposed to supplement the traditional experimental observation for accelerating advancements in related research^8–11^.

Machine learning models are a class of artificial intelligence (AI) for predicting peptide binding sites on proteins. Take the work of SPRINT-Str^12^ as an example, the authors employed structural information of proteins as the model’s features and a Random Forest (RF)^13^ to find the regions where peptides may bind. In addition, an approach that was grounded in protein sequence data denoted as SPRINT^14^, employed a Support Vector Machine (SVM)^15^ to predict protein-peptide binding sites. Most recently, SPPPred^16^ has been proposed as an ensemble-based classifier for distinguishing binding residues, through a genetic programming algorithm and related features. However, the major drawbacks of these algorithms are their dependence on expert knowledge and the quality of feature engineering, which have a substantial impact on their predictive performance.

While many of the protocols have been developed with machine learning, more recent works transition to newer and more accurate deep learning models. Algorithms rooted in deep learning have been successfully applied to a variety of protein-peptide binding site prediction tasks based on their data-driven feature engineering, without additional requirements for biological experts. For instance, GraphPPepIS^17^ employed multi-layer graph neural networks (GNN) to concurrently predict interaction sites for both proteins and peptides. PepNN^18^ can simultaneously update the encodings of protein and peptide through reciprocal attention, thereby predicting peptide binding sites on a protein. Additionally, popular methods such as AlphaFold-Multimer^19^ can also implicitly reveal the binding sites between the target and binder. However, those methods usually require ligand information or feature construction to guarantee precise prediction, which may reduce their practical utility. To address this issue, we propose a novel deep learning architecture called the Geometric Attention-based networks for Peptide binding Sites identification (GAPS), without excessive demand for laborious manual feature construction and related peptide ligand information.

The binding to ligands of a residue in the protein depends heavily on the surrounding residue environment^20^, therefore we applied the geometric feature to gain that information. Compared to previous modeling methods, the atom-based geometric information makes our model granularity smaller, increasing the likelihood of capturing inherent biological information among amino acid residues, and it also ensures the model’s translation-invariance and rotation-equivariance. Additionally, the attention mechanism^21^ was introduced to our model architecture to effectively optimize representations. In our work, the training size of protein-peptide interaction datasets is limited, which may affect the performance of our model. To further improve the predictive performance, we pre-trained GAPS on the protein-protein dataset and fine-tuned it to the downstream protein-peptide task (transfer learning), taking into account the biological similarity between proteins and peptides.

Our testing on benchmark datasets demonstrated the state-of-the-art (SOTA) performance of GAPS in the protein-peptide binding sites prediction by the combination of those tricks. GAPS was also implemented in other downstream tasks like identifying binding sites on the apo structures of protein receptors and predicting peptide binding sites based on the predicted receptor structures. Furthermore, to the best of our knowledge, GAPS stands as the first deep learning-based model for the protein-cyclic peptide binding sites prediction. The comprehensive experiments demonstrate the high accuracy and generality of our model in various binding site prediction tasks.

## 2. Results and Discussion

### 2.1 Geometric attention-based networks for peptide binding sites identification

The integration of the attention mechanism and geometric information within deep learning network architectures has demonstrated promising performance across various tasks^22,23^. Additionally, with significant advancements in protein structure prediction models, acquiring a substantial volume of high-confidence score predictions has become relatively straightforward^24–26^. These factors collectively contributed to the development of the GAPS.

GAPS exclusively utilized the spatial structure of proteins as input, represented as point clouds centered around atoms. The geometric feature, derived from distances and relative displacement vectors between these points, ensures the model’s translation-invariance and rotation-equivariance while obtaining the distance nearest neighbors for each point in the clouds (Fig. 1A). Subsequently, this geometric feature, alongside atomic scalar and atomic vector features extracted from points, were fed into the deep learning framework as Fig. 1B. This process led the framework to yield confidence scores for the given amino acid residues, signifying its potential as the binding sites (Fig. 1C). The input features of the model first passed through the geometric attention module, which continuously updated the atomic scalar and atomic vector embeddings as Fig. 1D. These embeddings were then directed to the geometric pooling module, converting atomic-level embeddings into residue-level embeddings. These residue-level embeddings were further inputted into the cross-attention module, facilitating mutual updates between scalar and vector representations (Fig. 1E). Following this, they entered the transformer encoder module to further refine residue-level embeddings. Finally, the matrixes of these embeddings were fed into a multi-layer perceptron yielding the confidence scores (more detail see methods). It’s also worth noticing that GAPS can handle other peptide binding site tasks as Fig. 1F.

**Fig. 1.**
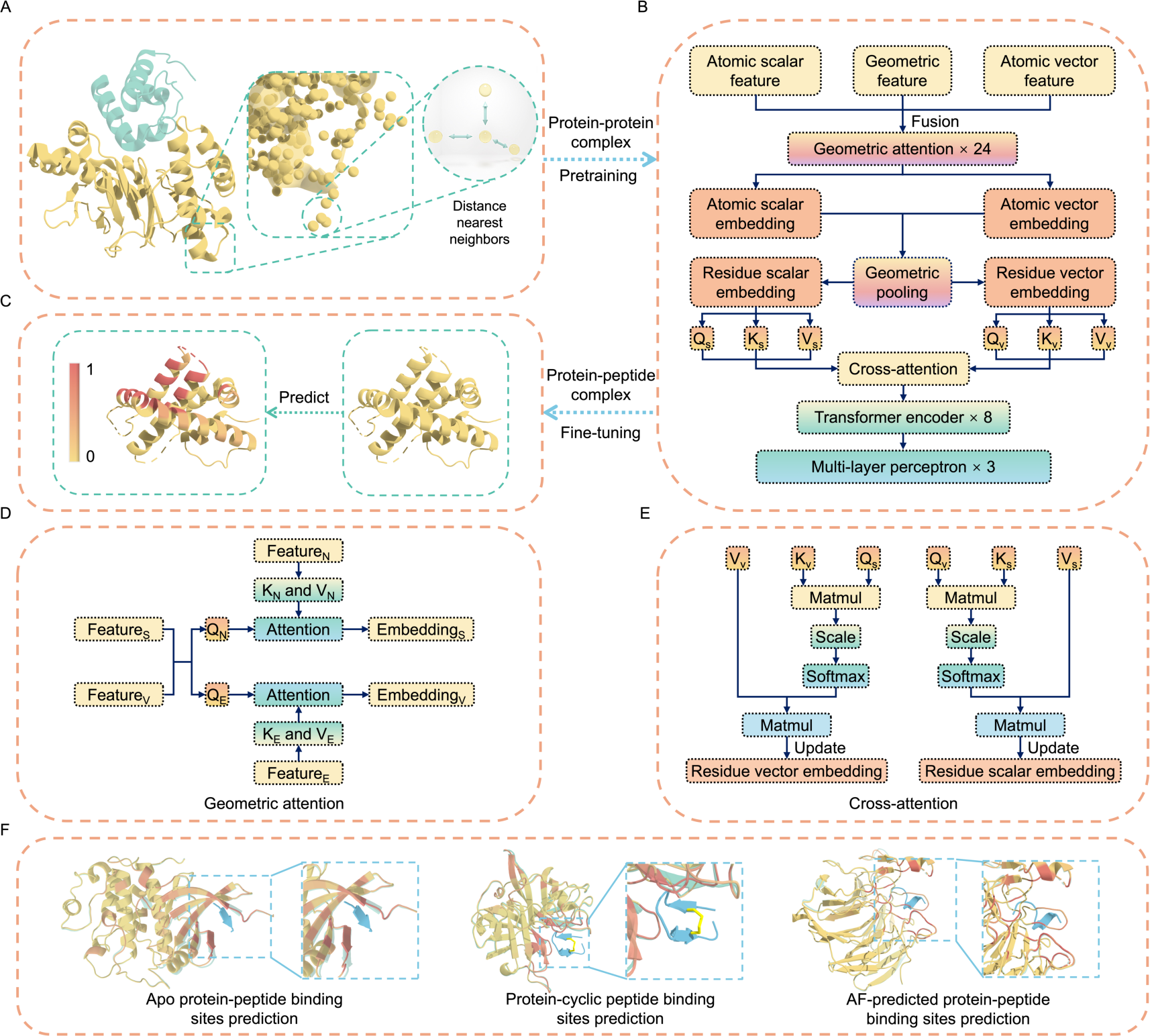
The overview of GAPS to predict the peptide binding sites. (A) The method to model the spatial structure of protein receptors. One chain within the protein-protein complex was colored in green to denote distinction. (B) The architecture of GAPS. Q_S_, K_S_, and V_S_ respectively represent the query, key, and value extracted from the residue scalar embedding. Q_V_, K_V_, and V_V_ respectively represent the query, key, and value extracted from the residue vector embedding. (C) The input and output of GAPS. Confidence scores of the prediction are represented with a gradient of color from yellow for non-binding residues to red for binding residues. (D) The architecture of geometric attention mechanism. Feature_S_, Feature_V_, Feature_N_, and Feature_E_ respectively represent the atomic scalar feature, atomic vector feature, distance nearest neighbors feature, and edge feature. Embedding_S_ and Embedding_V_ respectively represent the atomic scalar embedding and atomic vector embedding. (E) The architecture of cross-attention mechanism. (F) The various downstream tasks.

### 2.2 Protein-peptide binding sites prediction task

For the protein-peptide binding sites prediction, we first constructed a dataset named Data_finetuning_PepNN based on the dataset used by PepNN, a SOTA model for this task (see methods). Then, on this dataset, we adapt the pre-trained GAPS model that predicts protein-protein binding sites to predict protein-peptide binding sites. The resulting model can accurately predict protein-peptide binding sites as Fig. 2A. To evaluate GAPS’ performance, we conducted a comprehensive comparison on the test set, TS092. Compared to PepNN-Struct^18^, PepNN-Seq^18^, PBRpredict-flexible^27^, PBRpredict-moderate^27^, and PBRpredict-strict^27^, GAPS exhibits significant improvement in both AUROC and MCC metrics (Fig. 2B). Specifically, GAPS outperforms PepNN-Struct by a substantial margin, achieving a 6.2% increase in AUROC and a remarkable 7.3% improvement in MCC (0.917 and 0.482 vs 0.855 and 0.409 for AUROC and MCC, respectively).

**Fig. 2.**
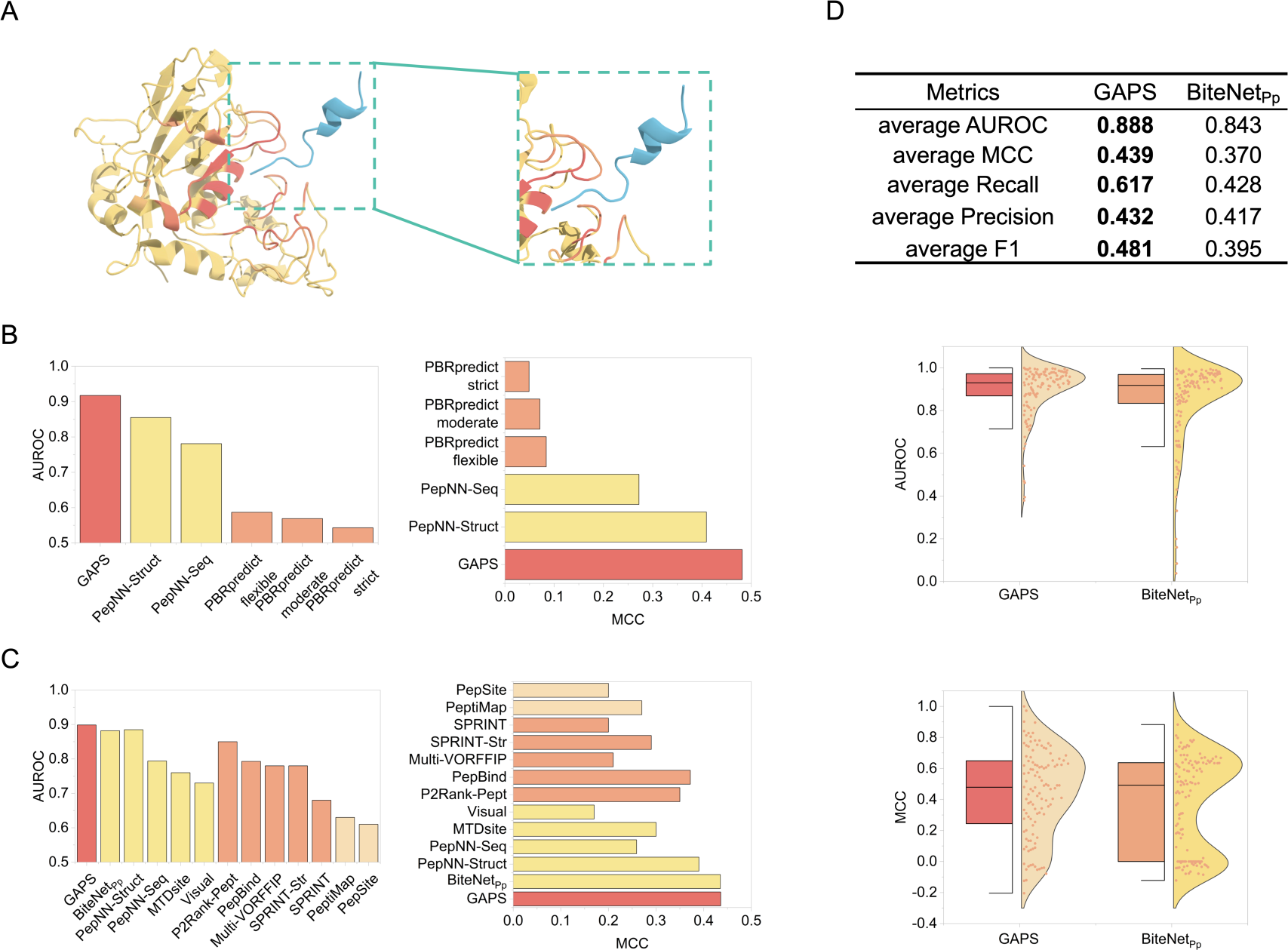
The predictive performance of GAPS in the protein-peptide binding sites task. (A) An example of predicted binding sites (PDB ID: 6R7W). The peptide colored in blue is added to show the quality of the prediction. (B) Comparison of GAPS and other methods on the TS092 with the AUROC and MCC metrics. (C) Comparison of GAPS and other methods on the TS125 with the AUROC and MCC metrics. GAPS is colored in red, deep learning-based methods are colored in yellow, machine learning-based methods are colored in orange, and others are colored in wheat. (D) Comparison between GAPS and BiteNet_Pp_ on the TS125 with average values of metrics. The better scores are shown in bold. The median is depicted by the center line of the boxplot, while the upper and lower edges delineate the interquartile range. The whiskers extend to 1.5 times the interquartile range. In the violin plot, the shaded areas portray the kernel probability density, indicating the proportion of samples. The results of methods other than GAPS are derived from the corresponding papers.

To further assess GAPS’ performance in predicting protein-peptide binding sites, we also fine-tuned the pre-trained GAPS using the Data_finetuning_BN and then compared it on the test set TS125 with other models including BiteNet_Pp_^28^, PepNN-Struct, PepNN-Seq, MTDsite^29^, Visual^30^, P2Rank-Pept^31^, PepBind^32^, Multi-VORFFIP^33^, SPRINT-Str^12^, SPRINT^14^, PeptiMap^34^, and PepSite^35^ (see methods). The results indicate that GAPS outperforms all other methods (Fig. 2C). Although GAPS did not surpass BiteNet_Pp_ by a large margin in terms of AUROC and MCC metrics (0.899 and 0.436 vs 0.882 and 0.435 for AUROC and MCC, respectively), it exhibits a substantial lead over BiteNet_Pp_ in terms of average AUROC and average MCC metrics as Fig. 2D. In practical applications, average AUROC and average MCC hold more significance than AUROC and MCC metrics, as they better reflect the predictive performance of individual samples. The diminished predictive capability of GAPS on the TS125 test set might be attributed to the small size of the Data_finetuning_BN which did not allow GAPS to reach its optimal performance due to inadequate training. These results show that GAPS demonstrates superior performance in predicting protein-peptide binding sites.

It’s worth noting that GAPS can accurately predict binding sites when there are multiple binding sites on a protein. For example, the Ku70/Ku80 heterodimer is a principal constituent of the DNA repair complex, primarily accountable for non-homologous end joining (NHEJ), forming a complex at both ends of DNA double-strand breaks. Each subunit of the heterodimer encompasses a von Willebrand factor A-like domain (vWA), facilitating interactions with other repair factors^36^. The Ku80 von Willebrand domain harbors two distinct peptide binding sites, each engaging with protein aprataxin and PNKP-like factor (APLF) and XLF, which can be precisely discerned by GPAS (Fig. 3A). GAPS also provided support for the prediction of protein-peptide binding sites on the multimer as shown in Fig. 3B. Overall, GAPS can make accurate predictions when the protein receptor’s PDB files are available. This underscores GAPS’ potential to contribute to the precise prediction of protein-peptide binding sites under diverse conditions, thereby aiding in the molecular-level elucidation of certain biological processes. In contrast, methods based on sequence and certain GNN or convolutional neural networks (CNN) approaches cannot deal with such scenarios.

**Fig. 3.**
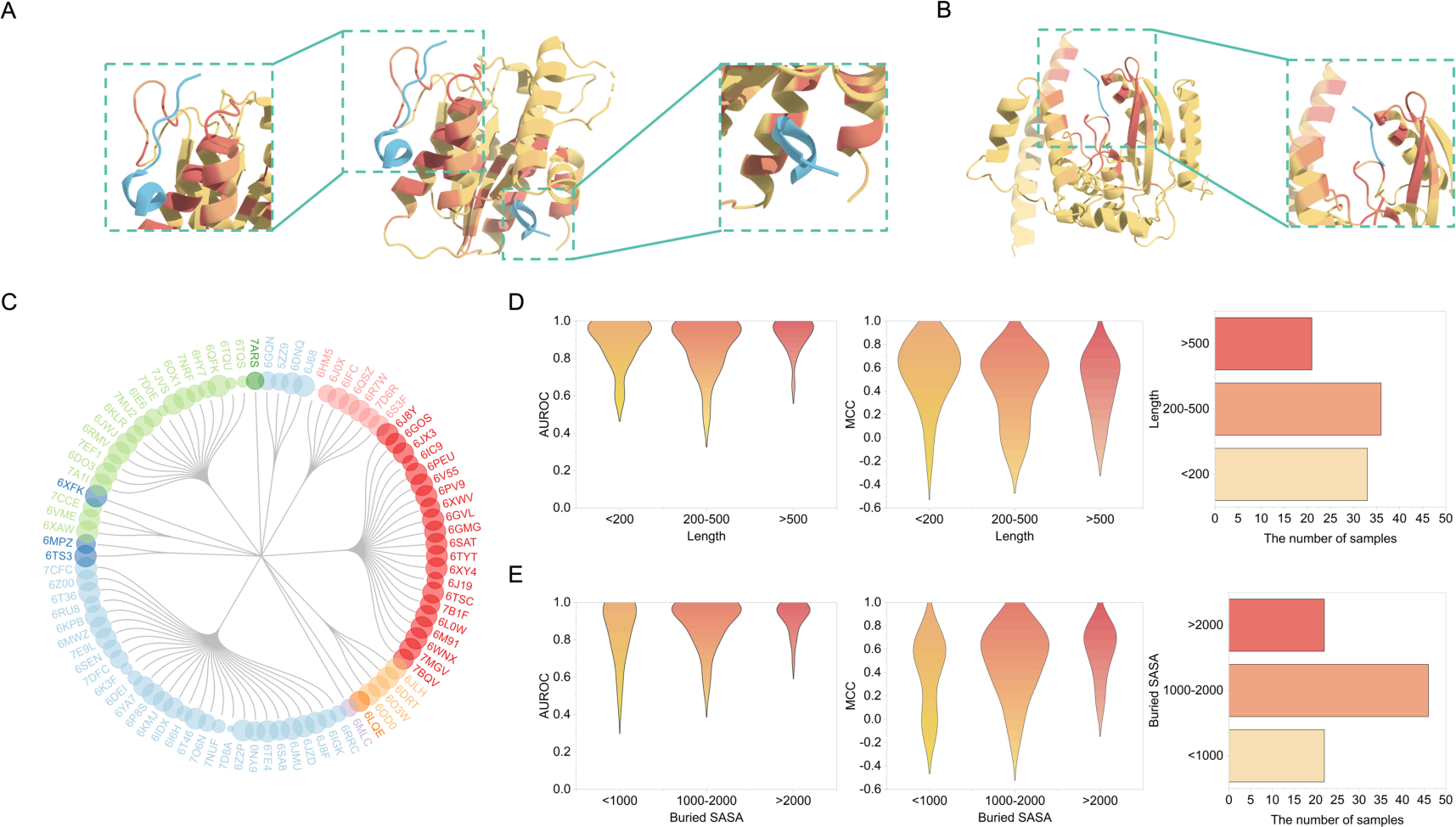
The robustness of GAPS. (A) An example of predicted multiple binding sites on a protein (PDB ID: 6TYT). The peptides colored in blue are added to show the quality of the prediction. (B) An example of predicted binding sites on the multimer (PDB ID: 6JZD). One chain is more transparent to distinguish, and the peptide colored in blue was added to show the quality of the prediction. (C) The clustering results on the TS092 using the Foldseek cluster. The size of the circle represents the AUROC score. (D) Distributions of AUROC and MCC predicted by GAPS correspond to the residue lengths of protein receptors on the TS092. (E) Distributions of AUROC and MCC predicted by GAPS correspond to the buried solvent-accessible surface area on the TS092.

To explore if GAPS exhibits any bias, we first employed the Foldseek cluster^37^, a structural-alignment-based clustering algorithm, to cluster the protein receptor structures in TS092 (see methods). The goal is to investigate whether GAPS favors specific structures. The results, depicted in Figure 3C, revealed that TS092 was grouped into 12 clusters by Foldseek cluster. Notably, structures 6LQE, 6MLC, 6XFK, and 7ARS formed individual clusters. The size of the circle represents the AUROC of GAPS predictions for that structure, where larger circles indicate higher AUROC values. This analysis demonstrated that GAPS does not exhibit a bias toward specific protein receptor structures. Furthermore, we explored the influence of the number of amino acid residues in the protein receptor on GAPS’ predictive performance, as shown in Figure 2D. TS092 protein receptors were divided into three groups based on the number of residues they contained. The results suggest that GAPS’ performance is not biased towards protein receptors with a specific number of residues. Finally, we investigated the impact of buried solvent-accessible surface area (buried SASA) on GAPS’ predictive performance (Fig. 3E). The findings revealed that GAPS’ performance is also not biased towards specific buried SASA values. Overall, these results underscore GAPS as a robust model for predicting protein-peptide binding sites, demonstrating its resilience to biases related to protein receptor structures, the number of residues, and the buried SASA.

### 2.3 Ablation experiments

To demonstrate the effectiveness of the pretraining and various modules within GAPS, we conducted an ablation study by removing the pretraining, transformer encoder, and cross-attention modules separately. We then impartially compared their performances on the TS092 test set as Fig. 4A.

**Fig. 4.**
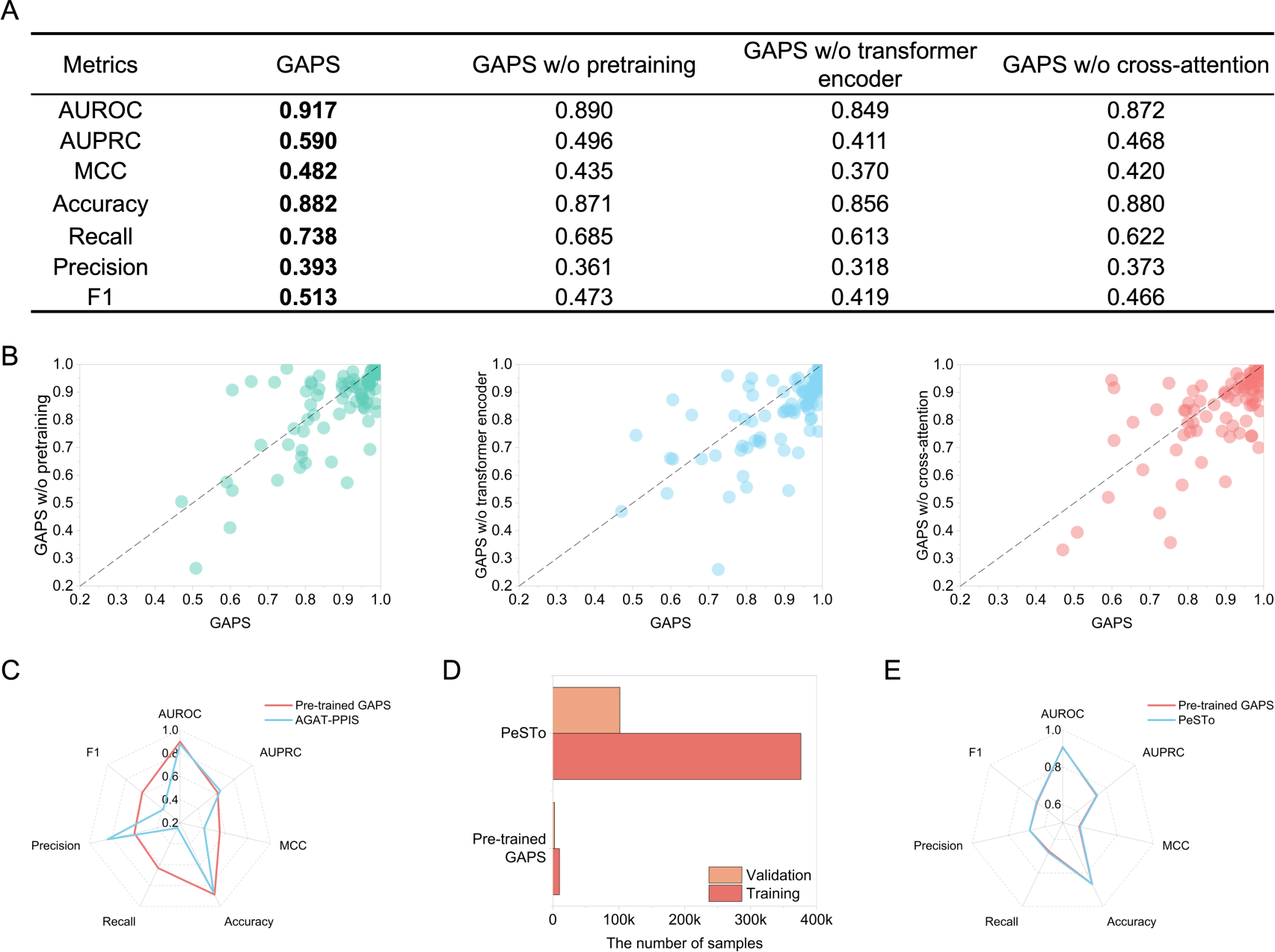
Ablation study. (A) Predictive performance comparison between GAPS and various ablation models on the TS092. w/o stands for without. The best scores are shown in bold. Neither GAPS w/o transformer encoder nor GAPS w/o cross-attention underwent pretraining. (B) AUROC comparison of GAPS and various ablation models for each sample within TS092. (C) Comparison of performance between pre-trained GAPS and AGAT-PPIS in the protein-protein binding sites prediction task. (D) Comparison of the size of training and validation sets used by pre-trained GAPS and PeSTo. (E) Comparison of performance between pre-trained GAPS and PeSTo in the protein-protein binding sites prediction task.

To investigate the necessity of pretraining, a model was trained from scratch directly on the Data_finetuning_PepNN. Comparing the performance of GAPS with and without pretraining demonstrates a substantial improvement when transfer learning is employed. This result emphasizes the value of transfer learning in enhancing the model’s capabilities for the protein-peptide binding sites prediction. We also observed that even without pretraining, GAPS’ performance surpasses that of the SOTA model, PepNN-Struct, with a 3.5% increase in AUROC and a 2.6% increase in MCC (0.890 and 0.435 vs 0.855 and 0.409 for AUROC and MCC, respectively). Protein-protein binding shares certain similarities with peptide-protein binding, and within the RCSB Protein Data Bank (PDB)^38,39^, there is a considerably larger amount of data available for protein-protein complexes compared to protein-peptide complexes. Therefore, including transfer learning can enhance GAPS’ performance on the protein-peptide binding sites prediction. Transfer learning also can provide superior initial parameters, thereby aiding the model converging more swiftly during the training process, steering clear of local minima and overfitting.

Next, we removed the transformer encoder module while retaining the cross-attention module, identically training from scratch directly on Data_finetuning_PepNN. As a result of removing the transformer encoder module, we observed a decrease in performance across all seven metrics. Finally, when the cross-attention module was removed while keeping the transformer encoder module, we found a significant decrease in the recall score. The higher recall score can indicate that better at identifying true positive binding residues in protein-peptide interaction prediction. In this context, a higher recall score means that the cross-attention module is better at capturing more of the actual binding residues, which can be particularly important when the goal is to avoid missing potential binding sites. Figure 4B displays the distribution of AUROC for each sample in TS092 across both GAPS and the ablated models, and we noticed that in most samples, GAPS outperformed the ablated models. In summary, the ablation study demonstrates the necessity of pretraining and various components within GAPS.

To assess the performance of pre-trained GAPS in predicting protein-protein binding sites, we compared it to the current SOTA models AGAT-PPIS^40^ and PeSTo^41^ (see Methods). Compared to the AGAT-PPIS model, the pre-trained GAPS model exhibits slightly lower AUPRC and precision scores. However, it significantly outperforms AGAT-PPIS in other metrics, with a 2.1% increase in AUROC, a 13.8% increase in MCC, a 1.9% increase in accuracy, a 38.3% increase in recall, and a 23.3% increase in F1 score (Fig. 4C). Furthermore, GAPS does not require the complex process of manual feature engineering used in AGAT-PPIS, making it a more streamlined and efficient model. Compared to the PeSTo model, despite the significantly smaller training and validation data used by the pre-trained GAPS model as shown in Fig. 4D, the differences between these two models across all seven metrics in protein-protein binding sites prediction task are negligible (Fig. 4E). These results indicate that the pre-trained GAPS outperforms AGAT-PPIS but performs comparably to PeSTo, highlighting the significant advantages of the GAPS architecture and showcasing its potential for application in other related tasks.

### 2.4 Apo protein-peptide binding sites prediction

In practical scenarios, we can only search binding sites on protein conformation without binding the ligand (apo structure). To assess GAPS’ performance in locating binding sites on apo forms, we constructed an independent test set Testset_apo based on data of HPEPDOCK^42^, which includes one-to-one corresponding apo and holo forms (see method). Prediction demonstrates that GAPS fine-tuned on Data_finetuning_PepNN can effectively identify binding sites on apo structures (Fig. 5A). Using GAPS for prediction in both holo and apo structures, we discovered that while GAPS performs better on holo structures compared to apo structures, its performance on apo structures is still remarkable, with AUROC of 0.842 (Fig. 5B). As shown in Fig. 5C, in the test set Testset_apo, there are several samples with relatively low AUROC scores in apo forms. To investigate whether the Root-Mean-Square Deviation (RMSD) between apo structures and holo structures affects GAPS’ predictive performance on protein-peptide binding sites, we divided the samples in Testset_apo into two groups based on the RMSD. We found that as the RMSD increases between the holo and apo forms, the accuracy of GAPS decreases (Fig. 5D). Perhaps, by employing molecular dynamics simulation tools on apo forms, we can further enhance GAPS’ predictive performance on these structures. These results indicate that GAPS can accurately predict peptide binding sites on apo protein receptors.

**Fig. 5.**
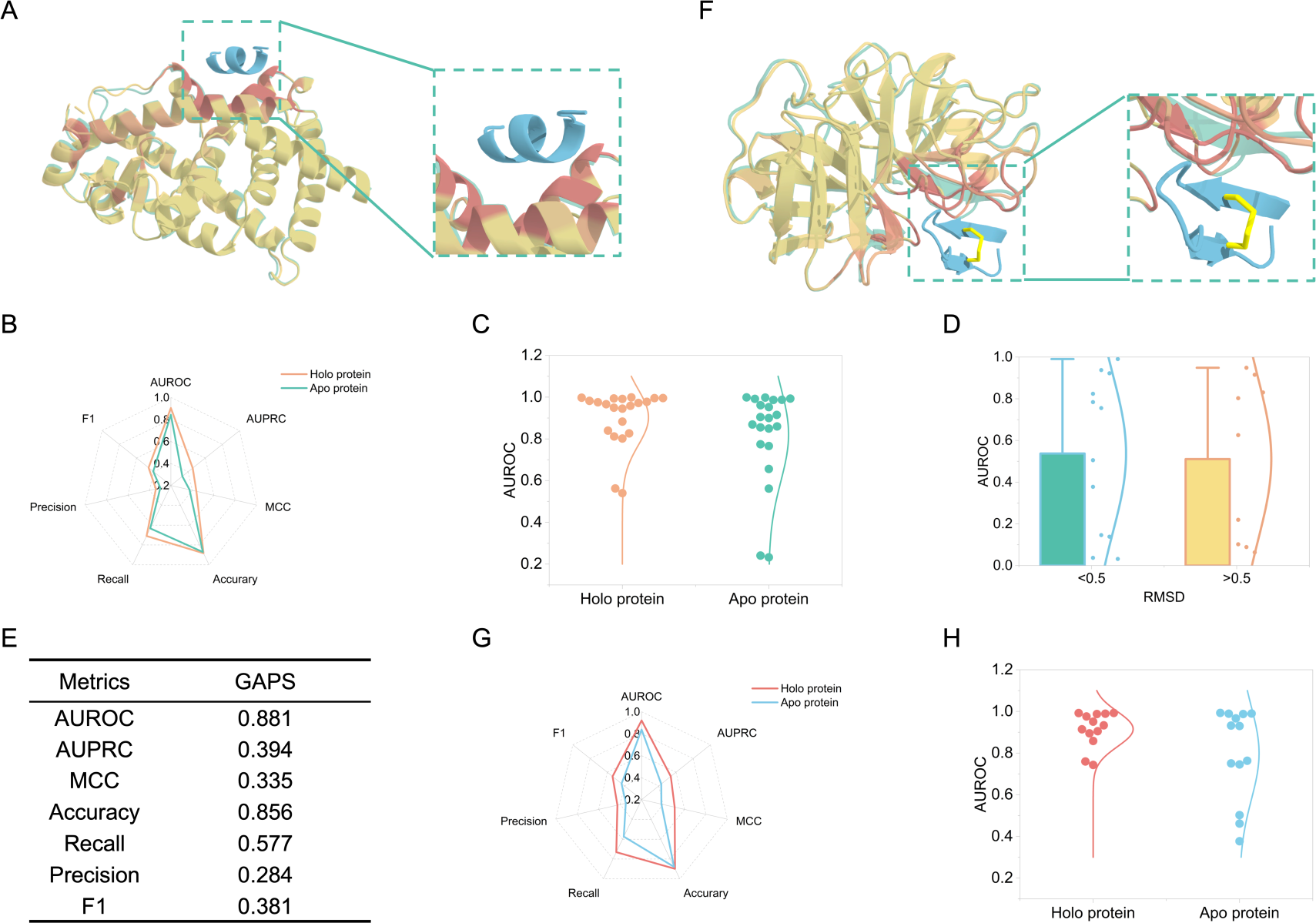
Apo protein-peptide and protein-cyclic peptide binding sites prediction. (A) An example of predicted binding sites on the apo proteins (PDB ID: 2HWQ). The peptide colored in blue and the holo protein colored in green based on the structure of the complex (PDB ID: 2FVJ) are added to show the quality of the prediction. (B) Comparison of GAPS’ predictive performance between apo protein and holo protein. (C) AUROC of GAPS for paired apo protein and holo protein on the Testset_apo. (D) Comparison of AUROC predicted by GAPS corresponding to the RMSD between apo protein and holo protein on the Testset_apo. (E) Predictive performance of GAPS in protein-cyclic peptide binding sites task. (F) An example of predicted binding sites on the apo protein for cyclic peptide (PDB ID: 4KGA). The cyclic peptide colored in blue and the holo protein colored in green based on the structure of the complex (PDB ID: 4K1E) are added to show the quality of the prediction. (G) Comparison of GAPS’ predictive performance between apo protein and holo protein for cyclic peptide. (H) AUROC of GAPS for paired apo protein and holo protein for cyclic peptide on the Testset_cyclic_apo.

### 2.5 Protein-cyclic peptide binding sites prediction

Due to their excellent membrane permeability, low toxicity, ease of synthesis, and high specificity, cyclic peptides have become robust candidates in the field of new drug development^43^. Accurately identifying the binding sites of protein-cyclic peptide complexes will significantly advance the development of cyclic peptide drugs. However, because of the scarcity of data on protein-cyclic peptide complexes, training a deep learning model from scratch for precisely identifying protein-cyclic peptide binding sites is impractical. Moreover, there is some commonality between protein-cyclic peptide and protein-peptide binding sites. Therefore, we have explored whether GAPS, the protein-peptide binding sites prediction model, can accurately predict protein-cyclic peptide binding sites using zero-shot learning.

Firstly, we built a protein-cyclic peptide binding sites prediction test set Testset_cyclic based on the data of protein-cyclic peptide complexes used in ADCP^44^ and HPEPDOCK2.0^45^ (see method). Subsequently, we employed GAPS to make predictions on this test dataset. The results demonstrate that GAPS can accurately predict protein-cyclic peptide binding sites. While GAPS exhibits lower performance in the task of predicting protein-cyclic peptide binding sites compared to its performance in predicting protein-peptide binding sites, its accuracy remains acceptable, with the AUROC of 0.881 on this test set (Fig. 5E).

To further evaluate GAPS’ performance in identifying protein-cyclic peptide binding sites on protein receptors without binding cyclic peptide ligands (apo structures), we refined the obtained test set Testset_cyclic for protein-cyclic peptide binding sites, retaining only the data with apo forms and called it Testset_cyclic_apo (see method). We used GAPS for prediction on apo structures, as depicted in Fig. 5F. Although GAPS’ performance on apo forms is notably lower than on holo forms, it still achieves the AUROC of 0.837. Additionally, to our knowledge, GAPS is the first deep-learning model currently supporting the prediction of protein-cyclic peptide binding sites. We also observed a greater disparity in predictive performance between holo structures and apo structures for cyclic peptide ligands compared to linear peptide ligands (Fig. 5G, 5H). This discrepancy might stem from inherent differences in the binding mechanisms between cyclic and linear peptides with the protein receptor, they are not entirely identical. We believe that with the accumulation of protein-cyclic peptide complex crystal structures, fine-tuning GAPS using such data will enhance its performance in predicting protein-cyclic peptide binding sites.

### 2.6 AF-predicted protein-peptide binding sites prediction

Acquiring crystal structures of proteins through experiments is challenging. However, with the development of protein structure prediction models, we can easily obtain many high-confidence predicted protein structures directly from amino acid sequences. To assess whether GAPS can utilize protein structures predicted by AlphaFold^46^ for accurate protein-peptide binding sites prediction, we selected 20 samples from the test set TS092, with release dates later than the date AlphaFold pulled training data. We then obtained the predicted structures of their protein receptors from the AlphaFold Protein Structure Database^47^ (see methods). Subsequently, we used GAPS to predict protein-peptide binding sites on these predicted structures, as illustrated in Fig. 6A.

**Fig. 6.**
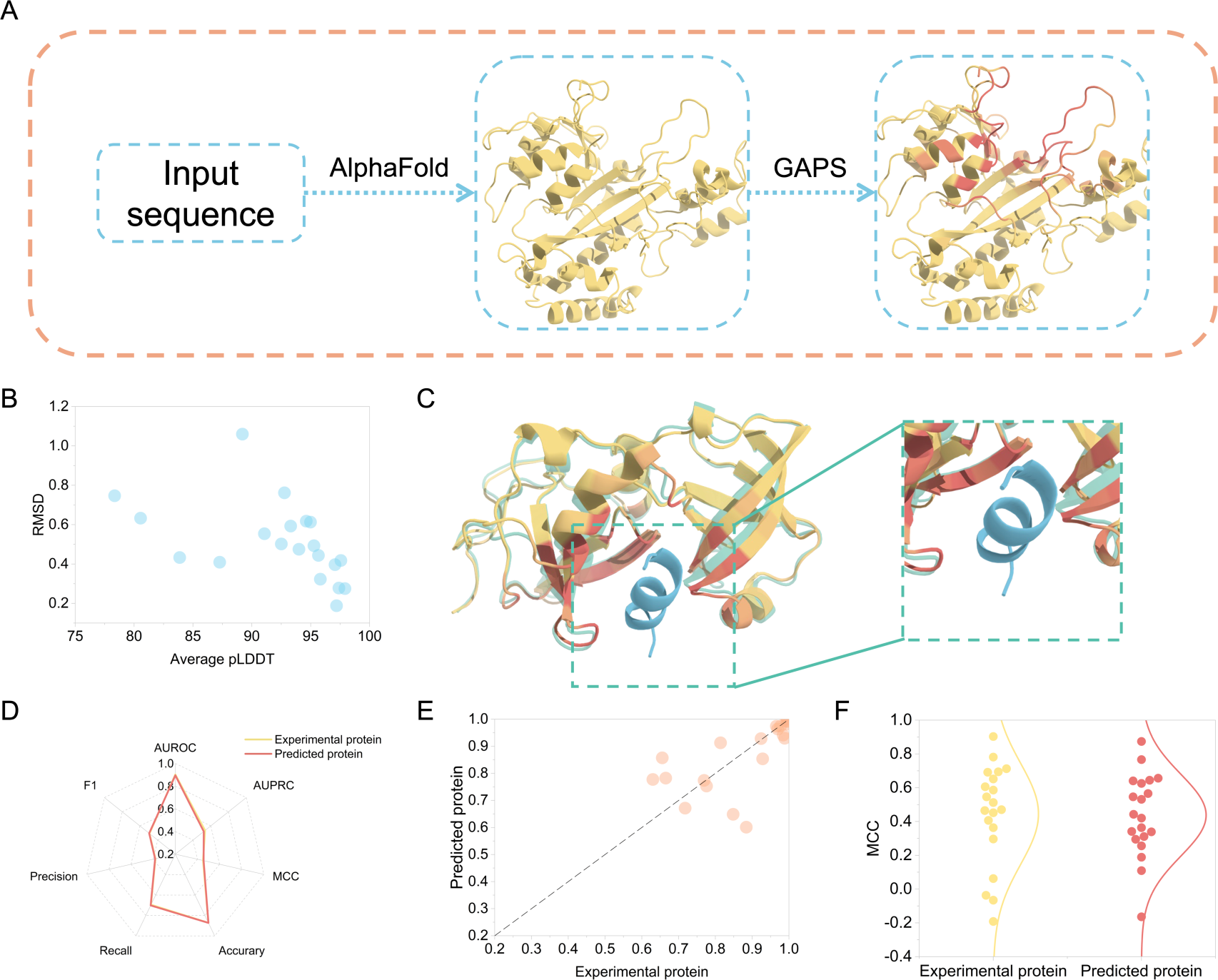
AF-predicted protein-peptide binding sites prediction. (A) The workflow for predicting the binding sites using GAPS based on predicted proteins. (B) The correlation between RMSD and average pLDDT. (C) An example of predicted binding sites on the proteins modeled by AlphaFold. The peptide colored in blue and the crystal protein colored in green based on the structure of the complex (PDB ID: 7CFC) are added to show the quality of the prediction. (D) Comparison of GAPS’ predictive performance between experimental protein and predicted protein. (E) AUROC comparison of experimental protein and predicted protein for each sample using GAPS. (F) AUROC of GAPS for paired experimental protein and predicted protein on the Testset_AFpredicted.

Analyzing the RMSD between predicted and experimental structures revealed that AlphaFold can accurately predict the majority of holo protein receptor structures, with only one predicted holo structure having an RMSD greater than 1 compared to the experimental holo structure. The linear relationship between average pLDDT and RMSD indicates that we can approximately assess the accuracy of the predicted holo structure based on the average pLDDT (Fig. 6B). As depicted in Figure 6C, GAPS can accurately predict protein-peptide binding sites on protein receptors predicted by AlphaFold. We have also observed that employing AlphaFold-predicted protein receptor structures for protein-peptide binding sites prediction does not impact the performance of GAPS (Fig. 6D, 6E, 6F). This might be correlated with AlphaFold’s extensive utilization of holo structures during training. In a word, using the predicted protein structures allows accurate prediction of protein-peptide binding sites, further expanding the applicability of GAPS.

## 3. Conclusion

Information regarding protein-peptide binding sites is important for understanding the protein-peptide interaction mechanism. With the advent of computer technology, many approaches have been proposed to predict the protein-peptide binding sites accurately. However, most of them necessitate expert knowledge or careful feature construction, which constrains their applicability in real-world scenarios. In this work, we have presented a deep learning model called GAPS for the prediction of protein-peptide binding sites by integrating atomic geometric feature extraction, the combined attention mechanism, and the transfer learning strategy. GAPS correctly predicted protein-peptide binding sites and achieved the SOTA performance for all metrics on various test sets. Through an ablation experiment, it is revealed that the pre-trained GAPS can achieve competitive performance in the protein-protein binding sites prediction task. Further analysis concerning different structure classes, the number of residues, and the buried SASA indicated that GAPS can gain a comprehensive performance on various protein receptors. In addition, the GAPS is used for the apo protein-peptide, the protein-cyclic peptide, and the predicted protein-peptide binding sites predictions. These extended experiments highlight the utility and generality of our model. We hope that this model can accelerate the development of related proteins and peptides research as an effective supplementary method.

## 4. Methods

### 4.1 Pre-processing

We only downloaded the first biological assembly from the PDB database and used it for training and if a PDB file contains multiple states, only the first is retained. All water molecules, hydrogen atoms, deuterium atoms, and heteroatoms are removed from this structure. When constructing a dataset for protein-protein binding sites, the PDB structures are split into independent target and ligand structures, according to the chain name. In the case of building a dataset for protein-peptide binding sites, structures with heavy atoms less than 255 are labeled as peptides, and the remaining parts of structures are considered protein targets. Within those data, the sets of residues on protein that are within 5Å (angstrom) of ligands are defined as binding sites. Protein targets with amino acid residue counts less than 48 or heavy atom counts exceeding 8196 are excluded from training.

### 4.2 Dataset

#### 4.2.1 Protein-protein binding sites pretraining data

The pretraining dataset for protein-protein binding sites prediction came from the work of ScanNet^48^, and we named this data Data_pretrain. The training, validation, and test subsets of this dataset comprised 10090, 2594, and 3316 samples, respectively. To fairly compare the performance of the GAPS pre-trained model in predicting protein-protein binding sites, we took the intersection of the Data_pretrain test set with the PeSTo test set, resulting in 526 samples, and we named this dataset Testset_PeSTo. Because the intersection of the Data_pretrain test set with the AGAT test set is too small, we took the difference of set between the Test_60 of AGAT-PPIS and the training and validation sets of Data_pretrain, yielding 29 samples, and named it Testset_AGAT-PPIS for further comparisons of pre-trained GAPS’ performance in predicting protein-protein binding sites.

#### 4.2.2 Protein-peptide binding sites fine-tuning data

The dataset for fine-tuning the pre-trained model to protein-peptide binding sites prediction came from the work of PepNN, and we named this data Data_finetuning_PepNN. The training, validation, and test subsets of this dataset respectively comprised 2373, 301, and 90 samples, and the test set is called TS092 (two protein receptors containing multiple peptide ligands were combined into one sample). To further assess our novel framework in predicting protein-peptide binding sites, we also used the BN_Pp_956 of BiteNet_Pp_ to fine-tune the pre-trained model and named it Data_finetuning_BN. The training, validation, and test subsets of this dataset respectively comprised 776, 119, and 116 samples, and the test set is called TS125 (eight samples were excluded from pre-processing and two samples were merged into a single sample due to being heterodimers).

#### 4.2.3 Other test data

We performed a subtraction between the holo data used by HPEPDOCK and the training and validation sets of Data_finetuning_PepNN. Subsequently, we extracted the apo data from HPEPDOCK. This process yielded 21 pairs of one-to-one corresponding samples. To facilitate a performance comparison of GAPS in holo and apo forms, we made certain modifications to these PDB files using PyMOL to ensure that their respective amino acid residue sequences were identical and named it Testset_apo.

The dataset for the protein-cyclic peptide binding sites prediction came from the work of ADCP and HPEPDOCK2.0, totaling 35 samples, and was named Testset_cyclic. In this dataset, we further filtered out samples that lacked apo structures. Finally, 13 paired apo-holo samples are retained, and we named it Testset_cyclic_apo.

We selected 20 samples from the test set of Data_finetuning_PepNN, with release dates later than April 30, 2018 (the time when AlphaFold pulled training data). Subsequently, we obtained the corresponding predicted structures using the AlphaFold Protein Structure Database. Finally, we obtained 20 paired samples and named them Testset_AFpredicted.

### 4.3 Representation

We model the spatial structure of proteins as point clouds. Each point in clouds includes a scalar state initialized as a one-hot encoding of its atomic element, while the vector state is initialized with zero. These scalar states, after passing through three fully connected layers, transform into the atomic scalar feature. Subsequently, they, along with the atomic vector feature formed by vector states and the geometric feature, are collectively input into the geometric attention module.

### 4.4 The architecture of GAPS

#### 4.4.1 Geometric attention

The geometric attention module consists of four sets of 24 layers. Each set includes 6 layers of geometric attention, with the number of distance nearest neighbors progressively increasing in the order of 8, 16, 32, and 64. Based on the geometric feature, the distance nearest neighbors feature and the edge feature can be represented. After concatenating the atomic scalar feature and atomic vector feature, we extract two queries from this merged result. Then, from both the distance nearest neighbors feature and edge feature, we extract a set of key and value pairs for each query. This process results in two sets of query, key, and value pairs. After that, the attention mechanism using those is performed to update the atomic scalar embedding and the atomic vector embedding. As the depth of the geometric attention layer increases, the number of distance nearest neighbors increases to expand the model’s receptive field. Its detailed calculation process is shown as follows:

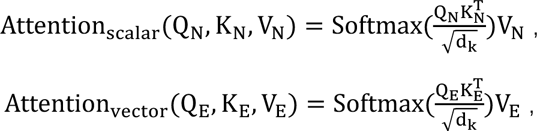

where Q_N_ and Q_E_ denote the two queries extracted from concatenated the atomic scalar feature and atomic vector feature. K_N_ and V_N_ represent the key and value extracted from the distance nearest neighbors feature, respectively. K_E_ and V_E_ represent the key and value extracted from the edge feature, respectively.

#### 4.4.2 Geometric pooling

Subsequently, the geometric pooling module takes the embeddings from the geometric attention module at the atomic level. It maps them to the residue level using a local multi-head mask, facilitating the pooling of atomic-level representation into residue-level representation. The canonical 20 amino acids are utilized to express the primary structure of the protein receptor. Each atom is segmented according to the amino acid residue to which it belongs, resulting in residue scalar embedding and residue vector embedding.

#### 4.4.3 Cross-attention

From the residue scalar embedding and residue vector embedding, a set of query, key, and value is extracted separately. These sets are then input into the cross-attention module. In this module, the query derived from residue scalar embedding utilizes the key and value obtained from residue vector embedding to execute the attention mechanism, thereby updating the residue vector embedding. Similarly, the cross-attention module employs the query from residue vector embedding alongside the key and value derived from residue scalar embedding to execute the attention mechanism, consequently updating the residue scalar embedding. Its detailed calculation process is shown as follows:

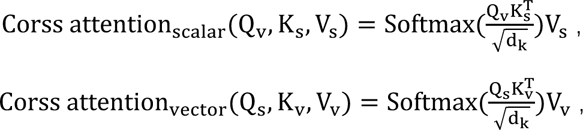

where the symbols Q_s_, K_s_, and V_s_ represent the query, key, and value extracted from residue scalar embedding, respectively. Meanwhile, Q_v_, K_v_, and V_v_ represent the query, key, and value extracted from residue vector embedding, respectively.

#### 4.4.4 Transformer encoder and multi-layer perceptron

The two sets of updated residue embedding are concatenated and input into an 8-layer transformer encoder module. Each layer has two sub-layers. The first sub-layer is to perform the multi-head self-attention mechanism, its detailed calculation process is shown as follows:

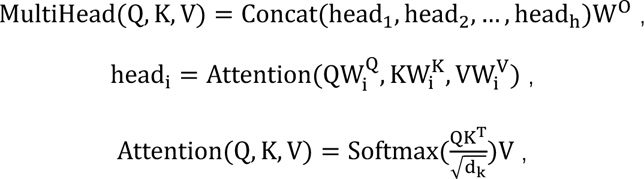

where the symbols Q, K, and V represent the query, key, and value extracted from concatenated residue embedding. All W denote the parameter matrices. Each layer consists of 8 heads. The second sub-layer is the position-wise fully connected feed-forward network, its detailed calculation process is shown as follows:

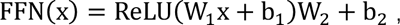

where W denotes the weight and b is bias. Each sub-layer is followed by residual connection and layer normalization. This module further optimizes the representations at the residue level on the protein. Finally, these representations are passed through a multi-layer perceptron with 3 layers to reduce them to 1 dimension, transforming them into a confidence score for the amino acid residue to be the binding site. Its detailed calculation process is shown as follows:

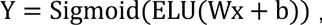

where W and b are the weight and bias, respectively.

### 4.5 Training

The model initially undergoes pre-training on the protein-protein binding sites dataset and is subsequently fine-tuned on the protein-peptide binding sites dataset. The model is implemented in PyTorch, and both the pre-training and fine-tuning phases employ the Adam optimizer with a learning rate of 1e-5. The loss function used is BCEWithLogitsLoss, and early stopping is based on the validation set’s loss to prevent over-fitting, with a maximum of 100 epochs for both pre-training and fine-tuning.

### 4.6 Evaluation

The GAPS pre-training model’s protein-protein binding sites prediction performance is compared with AGAT-PPIS and PeSTo, while the GAPS fine-tuning model’s protein-peptide binding sites prediction performance is compared with PepNN and BiteNet_Pp_. PeSTo is a SOTA protein-protein interfaces predictor based on the geometric transformer, and it can also accurately predict and distinguish binding interfaces involving nucleic acids, lipids, ions, and small molecules. AGAT-PPIS is a protein-protein interaction site predictor based on the graph attention network, and it is currently a SOTA model for predicting protein-protein interaction using graph-based methods. PepNN incorporates peptide ligand information to predict protein-peptide binding sites, achieving the highest performance in the protein-peptide binding sites prediction task. BiteNet_Pp_ leverages 3D CNN to predict protein-peptide binding sites by transforming protein structures into voxels.

Specifically, for the binding sites prediction task, we evaluate the model’s predictive performance by comparing the AUROC (Area Under the Receiver Operating Characteristic Curve), AUPRC (Area Under the Precision-Recall Curve), MCC (Matthews Correlation Coefficient), Accuracy, Recall, Precision, and F1 scores across the test set, its detailed definition as follows:

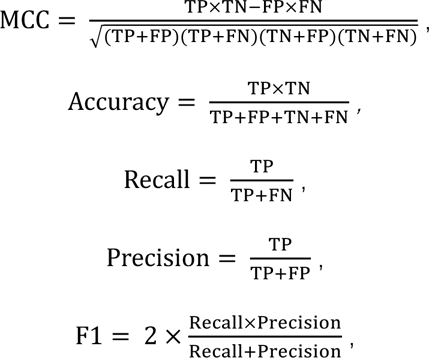

where TP (True Positive) represents the number of correctly predicted binding residues, TN (True Negative) represents the number of correctly predicted non-binding residues, FP (False Positive) represents the number of non-binding residues incorrectly predicted as binding residues, and FN (False Negative) represents the number of binding residues incorrectly predicted as non-binding residues. Exactly, if the model predicts a confidence score for an amino acid residue on the protein receptor that is equal to or greater than 0.5, it is classified as a binding residue. Conversely, if the predicted confidence score is less than 0.5, it is classified as a non-binding residue. We also compute AUROC, AUPRC, MCC, Accuracy, Recall, Precision, and F1 scores individually within each single PDB file. Averaging these metrics across all samples yields the average AUROC, average AUPRC, average MCC, average Accuracy, average Recall, average Precision, and average F1.

The RMSD is utilized to quantify the structural disparity between apo and holo structures or predicted and experimental structures. Its calculation formula is depicted as follows. We automatically computed this metric using the ‘align’ function within PyMOL.

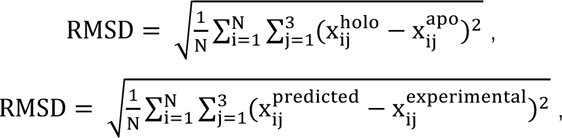

where N represents the number of amino acid residues in the protein receptor, 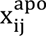 and 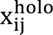 respectively represent the 3D coordinate values for the apo and holo structures and 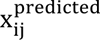 and 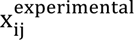 respectively represent the 3D coordinate values for the predicted and experimental structures.

The average pLDDT is obtained by averaging the pLDDT scores of each amino acid residue predicted by AlphaFold. The buried SASA is indirectly acquired through the ‘get_area’ function within PyMOL, and the calculation formula follows:

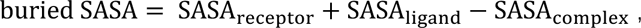

where SASA_receptor_ represents the SASA of the receptor, SASA_ligand_ represents the SASA of the ligand, and SASA_complex_ represents the SASA of the whole complex.

## 5. Acknowledgment

This project was supported by the Natural Science Foundation of Zhejiang Province (LD22H300004).

